# MEM-based pangenome indexing for *k*-mer queries

**DOI:** 10.1101/2024.05.20.595044

**Authors:** Stephen Hwang, Nathaniel K. Brown, Omar Y. Ahmed, Katharine M. Jenike, Sam Kovaka, Michael C. Schatz, Ben Langmead

## Abstract

Pangenomes are growing in number and size, thanks to the prevalence of high-quality long-read assemblies. However, current methods for studying sequence composition and conservation within pangenomes have limitations. Methods based on graph pangenomes require a computationally expensive multiple-alignment step, which can leave out some variation. Indexes based on *k*-mers and de Bruijn graphs are limited to answering questions at a specific substring length *k*. We present Maximal Exact Match Ordered (MEMO), a pangenome indexing method based on maximal exact matches (MEMs) between sequences. A single MEMO index can handle arbitrary-length queries over pangenomic windows. MEMO enables both queries that test *k*-mer presence/absence (membership queries) and that count the number of genomes containing *k*-mers in a window (conservation queries). MEMO’s index for a pangenome of 89 human autosomal haplotypes fits in 2.04 GB, 8.8*×* smaller than a comparable KMC3 index and 11.4*×* smaller than a PanKmer index. MEMO indexes can be made smaller by sacrificing some counting resolution, with our decile-resolution HPRC index reaching 0.67 GB. MEMO can conduct a conservation query for 31-mers over the human leukocyte antigen locus in 13.89 seconds, 2.5x faster than other approaches. MEMO’s small index size, lack of *k*-mer length dependence, and efficient queries make it a flexible tool for studying and visualizing substring conservation in pangenomes.

## 1 Introduction

There is a growing availability of pangenomes, including the Human Pangenome Reference Consortium (HPRC, n=94), the Vertebrate Genomes Project (VGP, n=16), and a recent pangenome for *Arabidopsis thaliana* (n=69) [27, 21, 16]. Pangenomes enable new ways of studying and visualizing variation, as well as the degree to which genomic sequences are conserved [24]. *K*-mers have proven to be a powerful tool for factoring, representing and indexing genomes. They have been used to power genome-wide association studies in plants such as barley and soybean [8, 13, 10], to identify single-copy genes in pangenomes [7, 8] and study sequence conservation [4].

However, indexing methods based on *k*-mers or de Bruijn Graphs require the value *k* to be set at index building time, limiting future queries to use that value of *k* only. PanKmer [4] supports only 31-mer queries, whereas length-31 substrings may not be the correct resolution at which to understand conservation across all genes or pangenomes [18]. Further, *k*-mer indexes can be large; e.g. an index consisting of a separate KMC3 database [11] for each haplotype in the HPRC requires 1.26 TB.

Alternative methods for indexing pangenomes also have drawbacks. Graph-based methods start with a computationally difficult reference-graph construction step. Accurate multiple alignments are difficult to create, requiring that some difficult repetitive sequences be masked first (e.g. the “dna-brnn” regions of the HPRC [27]).

Here we depart from the idea of a fixed length-*k* index by expanding on the notion of a “sequence landscape,” a vector of lengths of half-maximal exact matches between a query and reference text [6, 5, 23]. When computed between two sequences, a sequence landscape is equivalent to a vector of matching statistics (MS) lengths, from which maximal exact matches (MEMs) can be derived [22]. While MSs and MEMs have been applied successfully to classification [1, 2], they have not yet been used to *index* pangenomes.

We present a maximal exact match (MEM)-based compressed indexing approach called MEMO (Maximal Exact Match Ordered), along with new concepts enabling both lossless and lossy pangenome indexes. MEMO indexes MEMs between a “pivot” genome selected from the pangenome with respect to all the other genomes. MEMO can then answer any-length *k*-mer queries for *k*-mers drawn from the pivot.

MEMO builds from a few methodological principles. First, the MEMs indexed by MEMO are sufficient for answering any *k*-mer membership or conservation query for any *k* as long as the *k*-mer is from the pivot. Second: it is also sufficient to store only the intervals representing overlaps between consecutive MEMs, helping to reduce index size. Third: we also use a variation on MEMs called “order-MEMs”, obtained by re-sorting the values in the matching statistics vectors. Order-MEMs speed up conservation queries both by enabling early stopping, i.e. the ability to return a correct answer after examining some but not all of the orders, and by enabling lossy compression as described below.

We also introduce two ideas to further reduce index size with lossy compression. The first builds on the use of order-MEMs; once arranged as orders, we can discard some orders— and potentially a large fraction of the MEM intervals—as long as the user is satisfied with coarse-grained answers to conservation queries. A coarse-grained answer does not convey the exact number of genomes in which a substring occurs, but could instead return e.g. the largest percentage such that the substring occurs in at least 10%, 20%, …, 90% or 100% of the genomes (a “decile conservation query”). The second idea builds on the observation that if we limit the user to making *k*-mer queries where *k* is greater than a threshold length *t*, we can reduce the index size further by discarding MEMs with length ≤ *t*. If we operate using MEM overlaps rather than MEMs themselves, we can apply an inverted version of that principle; i.e. we can enable *k*-mer queries for *k* less than a threshold length *t* while discarding MEM overlaps greater than a threshold length.

Finally, we offer the practical insight that indexes over pangenomic MEMs—as well as the variants discussed here—are quite compressible, due both to the inherent repetitiveness of pangenomes, and to the inherent inefficiency of how offsets and lengths are stored in BED files. This compression is not exploited when intervals are simply placed in BED files, nor is it well exploited by standard compression approaches compressing BED files, such as gzip or tabix [14]. Approaches that use *columnar* compression such as Apache Parquet [26] are key to achieving the needed degree of compression while still enabling fast queries.

Here we test MEMO by comparing its index size and query speed to existing *k*-mer-based methods for membership and conservation queries. We find that MEMO consistently yields the smallest index size, sometimes orders of magnitude smaller than those from comparable approaches. We show how MEMO’s index scales well to large pangenomes, and that its lossy and lossless strategies for reducing index size were effective, ultimately fitting the HPRC index for coarse-grained conservation queries in less than 1 GB. Finally, we demonstrate MEMO’s utility in visualizing and exploring sequence conservation in a pangenome by visualizing sequence conservation in the region of the human genome containing the Human Leukocyte Antigen (HLA) genes.

## 2 Methods

### 2.1 Methods Overview

MEMO is a pangenome index enabling arbitrary-length *k*-mer membership and conservation queries. If the pangenome consists of *N* genome sequences, a *k*-mer membership query returns a length-*N* vector of true/false values indicating the presence/absence of the *k*-mer in each genome. A *k*-mer conservation query returns the number of genomes that the *k*-mer occurs in, which is an integer in [1, *N*]. Both membership and conservation queries are limited to *k*-mers that occur in a particular genome, called the “pivot.” Thus, a membership query will always return at least one “true” value (for the pivot) and the conservation query result will always return a value ≥ 1.

The index works by pre-computing and indexing the MEMs between one of the genomes (the “pivot”) and all the others. The user may pose queries using only *k*-mers from the pivot; i.e. the user specifies the interval within the pivot containing the query *k*-mers. The index answers such queries by examining whether and how the query intervals overlap the indexed MEM intervals. Some pangenomes will have a natural choice of pivot. For example, the T2T-CHM13 assembly is the most complete [19]. But this is also a limitation of the MEMO approach; when there is more than one natural pivot, the user may need to build multiple indexes for multiple choices of pivot.

### 2.2 Preparing for the MEMO index

To build the index, we first compute vectors of matching statistics (MSs), also called sequence landscapes [6], between the pivot and the other genomes (Figure 1A). This yields a matrix of MSs, where the rows are genomes and columns are pivot coordinates (Figure 1C, bottom). MEMs map one-to-one to peaks in the matrix, i.e. instances where one MS is not less than the MS to its right. Presence or absence of a *k*-mer from pivot offset *i* is determined by asking whether it falls entirely within a MEM. (Figure 1C, left middle). MEMO also implements a complementary approach, which considers whether a *k*-mer contains an overlap region between consecutive MEMs (Figure 1B) with overhang on either side, in which case it is not present.

**Figure 1.**
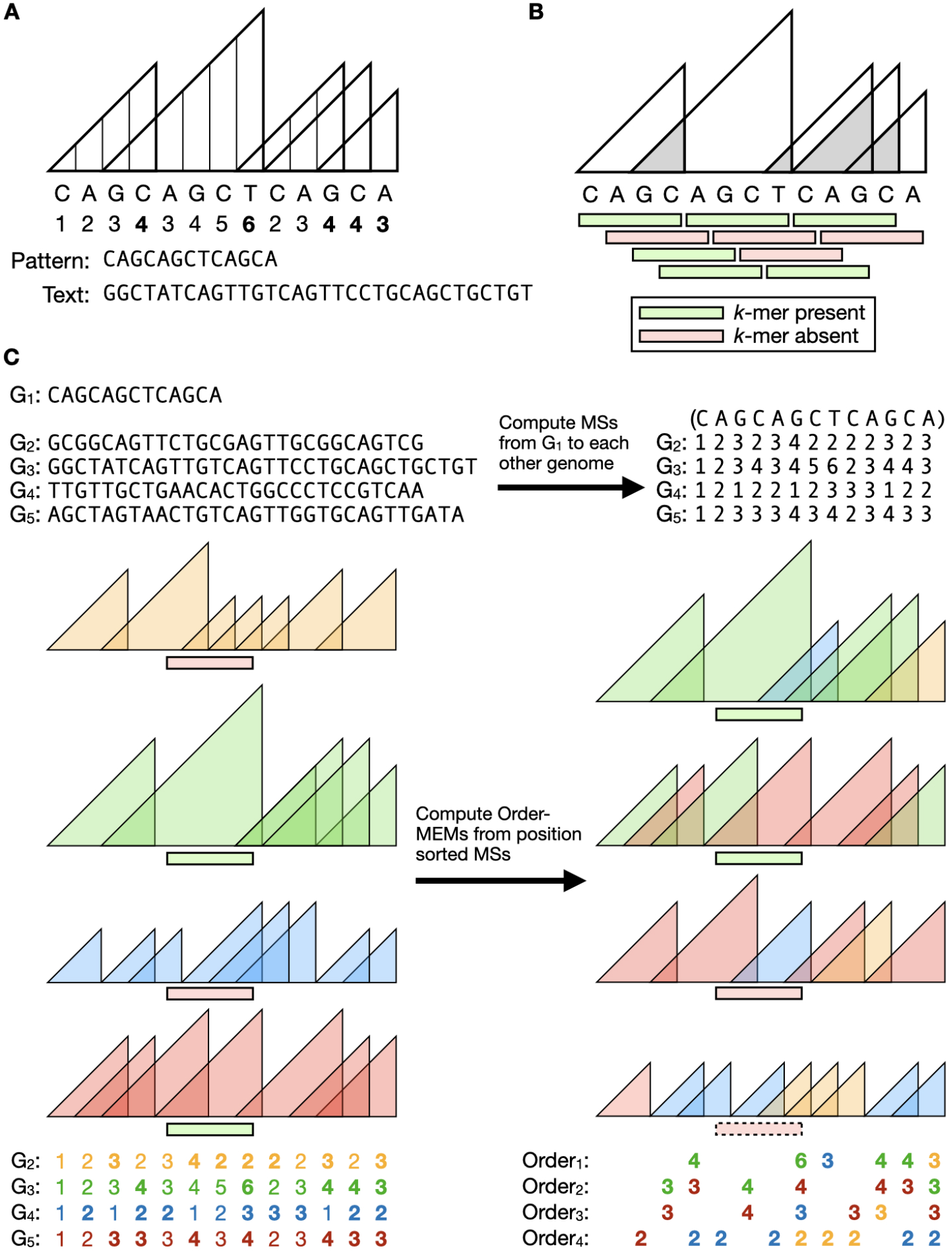
MEMO index outline. **A** Numbers below the pattern (i.e. pivot genome) are matching statistic (MS) lengths with respect to the Text (i.e. other genome). Triangles represent MS positions and lengths. MS peak lengths are bolded; these correspond to maximal exact matches (MEMs). **B** Presence/absence of the Pattern’s *k*-mers depicted as green and red rectangles. Grey triangles represent overlaps between consecutive MEMs. **C** Order-MEM creation from a pangenome. Top left: Sequences of anchor genome (*G*_1_) and genomes (*G*_2_-*G*_5_). Top right: MSs matrix of match lengths between *G*_1_ and *G*_2_-*G*_5_. Bottom: Order-MEMs found from MSs. MEM triangles and MSs of *G*_2_-*G*_5_ are colored individually. MEM lengths are bolded. Order-MEMs of a single order are composed of MEMs of varying genomes. *K*-mer presence/absence of an example query are depicted in green and red. The order-MEM *k*-mer query enables early stoppage, as depicted by the *k*-mer in the dotted outline.

To answer conservation queries, MEMO uses a rearranged version of MSs, whereby the MS matrix is first sorted along its column axis. After this, a row no longer represents MSs with respect to a particular genome, but instead represents “order” MSs with respect to the entire pangenome (Figure 1C, right). MEMs derived from the reorderd matrix are called order-MEMs. A *k*-mer fully contained in an order-MEM from order *x* occurs in at least *x* other genomes in the pangenome.

For all index types, MEM intervals are indexed as a columnar-compressed BED file, with an extra column containing an identifier for the genome of origin (for membership queries) or a number indicating the rank of the order statistic (for conservation queries).

### 2.3 Preparing for the MEMO index

A MEMO index is derived from an initial set of matching statistics (MS) vectors. MSs are half-maximal exact matches between a pattern *P* [1..*m*] and text *T* [1..*n*] that cannot be extended to the left without introducing a mismatch or reaching the end of a string (Figure 1A). We define the MSs of *P* as an array MS[1..*m*] where MS[*j*] is the length of the longest suffix of *P* [1..*j*] occurring in *T*. We note that this definition of MS is reversed with respect to how it is defined in some other work. This is because some algorithms for computing MS naturally work in right-to-left direction. For simplicity, we will define and discuss MS[1..*m*] as though it is computed left-to-right.

By definition, successive values of MS[1..*m*] have the *sawtooth property*:

#### ▶ Lemma 1.

MS[*j*] − MS[*j* − 1] ≤ 1, *j* ∈ (2, *m*]

On a collection of genomes *G* = [*G*_1_, *G*_2_, …, *G*_*t*_], MEMO factors the index building process into (*t* − 1)-pairwise comparisons between the pivot genome *P* = *G*_1_ and each of the other (*t* − 1)-genomes. MEMO uses MONI to compute these MS vectors [22]. Specifically, MEMO builds a MONI index over each genome and its reverse-complement sequence, appending $ to the end of each sequence in order to mark boundaries. MEMO then queries the pivot against each index to find MSs. These MSs are arranged in a *m* x *t* − 1 matrix. While MEMO uses MONI, any tool capable of finding all MEMs or matching statistics could be used, such as SPUMONI [1] or MUMmer [17].

### 2.3 MEMO index

#### 2.3.1 MEMO index with genome annotation

The full vector of matching statistics can be more concisely represented as a vector of MEMs. A MEM is an exact substring match between the genomes that cannot be extended left or right without introducing a mismatch or reaching the end of a genome. MEMs can be derived from MSs and vice versa: MS[*j*] is a MEM if and only if:

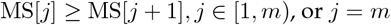

In other words, a one-to-one mapping exists between MEM and “peaks” of the sawtooth (Figure 1A), where the above expression defines a peak.

MEMO finds all MEMs between the pivot and each other genome in this way. Also, MEMO computes and stores the *overlaps* between consecutive MEMs, which we call “overlap MEMs.” These are illustrated in Figure 1B. Overlap MEMs are stored as zero-indexed, half-open intervals (i.e. with the low offset being inclusive and the high offset being exclusive) in a columnarly-compressed BED file. Overlaps between adjoining consecutive MEMs, where one MEM ends exactly where the next MEM starts, are stored as an interval with the same value for its start and end.

#### 2.3.2 MEMO index with order annotation

MEMO can also index and perform conservation queries with respect to “order-MEMs.” For the pangenome *G* = [*G*_1_, *G*_2_, …, *G*_*t*_], we define a matching-statistics matrix *L*[1..*t*][1..*m*] such that *L*[*i*] = MS[1..*m*] with respect to *T* = *G*_*i*_ and *P* = *G*_1_. We define the order-matching-statistics (order MS) matrix O[1..*t*][1..*m*] as the result of sorting *L* along its columns in descending order, such that O[1][*j*] ≥ O[2][*j*] ≥ … ≥ O[*t*][*j*] (Figure 1C, botom). Though the rows of O are not defined the same way as those of *L*, it is notable that order-MSs also have the sawtooth property:

##### ▶ Lemma 2.

O[*i*][*j*] − O[*i*][*j* − 1] ≤ 1 *for i* ∈ [1, *t*], *j* ∈ (1, *m*]

**Proof**. By sorted order of O, there can be at most *i* entries in O[1..*t*][*j* − 1] strictly greater than O[*i*][*j* − 1]. Let *π* be the permutation that sorted L[1..*t*][*j* − 1], such that O[*i*][*j* − 1] = L[*π*^−^1(*i*)][*j* − 1]. Then for any *i*^*′*^ *< i*, Lemma 1 ensures O[*i*^*′*^][*j* − 1] + 1 ≥ *L*[*π*^−1^(*i*)][*j*]. Since these values are then sorted in O[1..*t*][*j*] it follows that there are at most *i* entries in O[1..*t*][*j*] strictly greater than O[*i*][*j* − 1] + 1, guaranteeing that O[*i*][*j*] ≤ O[*i*][*j* − 1] + 1, and hence O[*i*][*j*] − *O*[*i*][*j* − 1] ≤ 1.

Similarly to how we use the 1-to-1 mapping between MS peaks and MEMs, we use this same mapping to extract “order-MEMs” from O. That is, O[*i*][*j*] is a peak if and only if: O[*i*][*j*] ≥ O[*i*][*j* + 1], *j* ∈ [1, *m*) or *j* = *m*. For conservation queries, MEMO computes and indexes order-MEMs. Similarly to how MEMO handles typical MEMs, MEMO indexes the overlap between consecutive order-MEMs rather than the order-MEMs themselves. These order-MEM overlaps are encoded as intervals in a BED file, similarly to 2.3.1, but with the interval’s annotation set to its order (i.e. row in the O matrix) instead of its genome ID.

### 2.4 Quantile-sampled conservation indexes

Conservation queries do not always need to be answered at full resolution. The HPRC pangenome includes around 90 human haplotypes; the difference in conservation level between a *k*-mer that occurs in exactly 71 genomes versus one that occurs in exactly 72 is not large, and may not be relevant to the scientific question. Also, such small relative differences would be hard to distinguish in a visualization.

For situations where a coarser resolution is sufficient, we propose a lossy-compression strategy called quantile sampling. Say that is sufficient for the user to learn whether a *k*-mer is present in at least *x*% of genomes, where *x* is a multiple of 10, i.e. count *deciles*. We subsample rows of the O matrix to include only those rows coinciding with movements into the next-highest decile. For instance, for the 89 haplotypes of the HPRC, we would sample rows corresponding to order-MEMs present in at least 9 samples, since that corresponds to order-MEMs present in at least 10% of samples. Likewise, we would sample the rows corresponding to order-MEMs present in at least 18, 27, …, 89 samples, since those represent order-MEMs present in at least 20%, 30%, …, 100% of samples. Rows of O not sampled in this way are discarded. Note that this scheme is easily adapted to any set of quantiles, e.g. quartiles, percentiles, etc.

#### 2.4.1 Columnar-compressed index

MEMO indexes are compressed and indexed using PyArrow, a Python API for Apache Arrow. The index is stored in an Apache Parquet file. A Parquet file is organized into chunks of rows (MEMs), where rows within a chunk are laid out and compressed in a columnar fashion, i.e. with data items arranged in column-major order. This yields a better compression ratio than if the data were indexed in row-major order, since it brings the values most likely to be redundant (e.g. MEM starting coordinates) into closer proximity.

Parquet supports efficient column-wise queries, compression, and decompression. Columns of a MEM index are compressed using the ZSTD codec. Rows are factored into blocks such that a single block occupies about 0.5 GB.

While other compression and indexing methods, such as bgzip and tabix [15], could be used instead of Parquet and ZSTD, we found this combination to provide excellent compression and speed in practice, as seen in the Results.

### 2.5 Queries

Say we have already computed the matching statistics for pivot genome *G*_1_ with respect to another genome *G*_2_. Given the matching statistics, we can collect the start and end coordinates of all of the MEMs, storing these in an interval-based data structure. By the definition of a MEM, we know that a *k*-mer *G*_1_[*i*..*i* + *k* − 1], is present in *G*_2_ if and only if and only if the interval [*i, i* + *k* − 1] is entirely contained in a MEM. This is true regardless of *k*; that is, we can answer arbitrary-length presence/absence queries for *G*_1_’s substrings with respect to *G*_2_ using only the MEM intervals.

Given an array of all the MEMs in order according to their starting coordinate, we can derive a second array of “overlap-MEMs.” Specifically, from a consecutive pair of MEMs [*i, i* + *𝓁* − 1], [*j, j* + *𝓁*^*′*^ − 1] where *j > i*, we derive a single overlap MEM [*j, i* + *𝓁* − 1]. We can perform presence/absence queries with respect to overlap MEMs:

#### ▶ Lemma 3.

*Consider a k-mer interval from G*_1_ [*i, i* + *k* − 1]. *If there exists an overlap-MEM interval* [*j, j* + *𝓁* − 1] *such that i < j and i* + *k > j* + *𝓁, then the k-mer is not present in G*_2_.

**Proof**. By definition of a MEM, we know that the *k*-mer at interval [*i, i* + *k* − 1] is present in *G*_2_ if and only if it is entirely contained within a MEM. Also by definition of a MEM, a MEM interval cannot contain another MEM interval.

If there exists an overlap-MEM [*j, j* + *𝓁* − 1] such that *i < j* and *i* + *k > j* + *𝓁*, then the *k*-mer interval [*i, i* + *k* − 1] is contained in neither the leftmost nor the rightmost of the two MEMs that created the overlap-MEM. Because these MEMs were consecutive, no other MEM exists that could span the *k*-mer interval.

Therefore, we have two distinct ways to test for the presence/absence of a *k*-mer from *G*_1_: (a) we can test whether the *k*-mer interval is entirely contained in a MEM, in which case it is present, or (b) we can test whether the *k*-mer interval spans an overlap-MEM including “overhang” on both sides, in which case it is not present. In practice, we generalize the query from pivot *G*_1_ to genome *G*_2_ to all (*t* − 1)-genomes (*G*_2_, …, *G*_*t*_) of the pangenome. A single index file contains all the MEM intervals—each interval annotated by document ID or order.

We apply the substring presence/absence query on overlap MEMs for the *k*-mer membership query:

#### ▶ Definition 4.

*For G*_1_[*i* : *j*] *and length k, yield G*_1_[*x* : *x* + *k* − 1] *in G*_*n*_ *for x* ∈ [*i, j* − *k*], *n* ∈ [2, *t*]

Given the overlap MEM intervals *M* of the membership index of *t* genomes, MEMO can compute arbitrary *k k*-mer presence/absence on any *l*-length substring of the pivot.

#### Algorithm 1

MEMO - Membership query

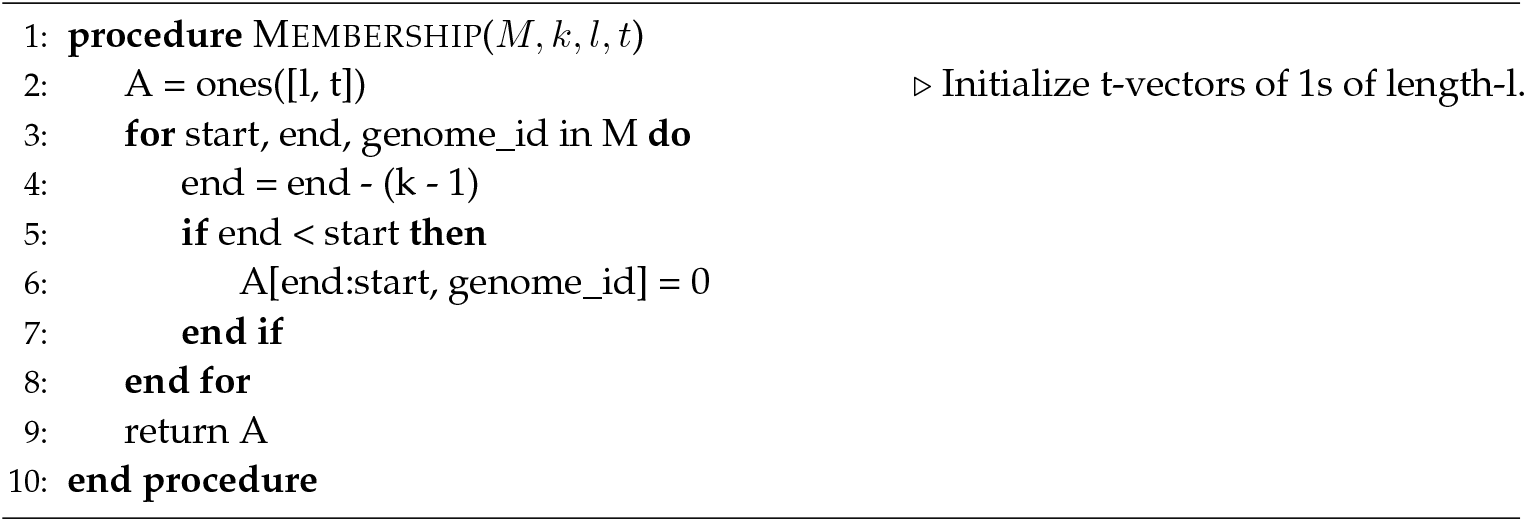

Likewise, MEMO applies the substring presence/absence query on order-MEMs for the conservation query problem:

#### ▶ Definition 5.

*For G*_1_[*i* : *j*] *and length k, yield 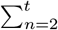*

MEMO uses a similar algorithm as its membership query (Algorithm 1), but outputs the last order a *k*-mer is present in.

#### Algorithm 2

MEMO - Conservation query

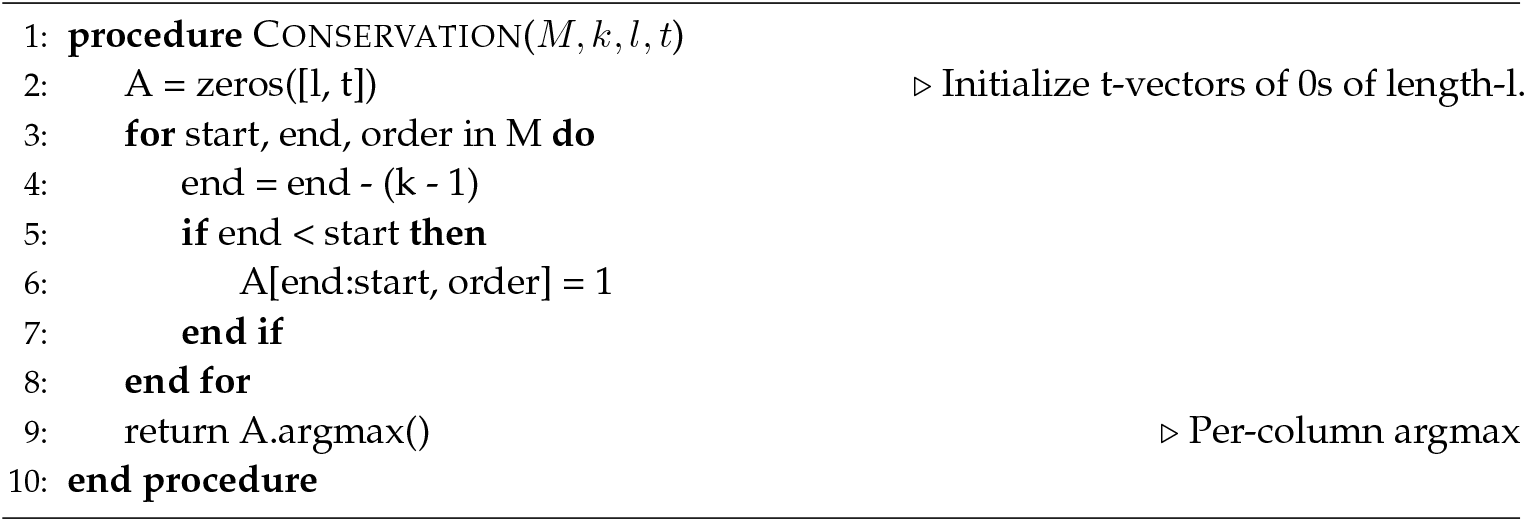

### 2.6 KMC3 index and query

We use KMC3 as a comparison to MEMO’s membership and conservation queries [11]. For the KMC3 membership index, we created a KMC3 database for each genome in the pangenome with the count of each canonical *k*-mer present transformed to 1. The KMC3 membership query uses `samtools faidx` to isolate the query substring from the query FASTA and KMC3 API’s `GetCountersForRead` function to query each *k*-mer in the substring against each KMC3 database.

The KMC3 database for the conservation query is constructed by taking the union/sum of each of the genome-specific KMC3 databases. That is, the count associated with a *k*-mer in the joined database is the sum of presence/absence values in each genome. This straightforwardly provides answers to conservation queries.

## 3 Results

We compared MEMO index sizes and query speeds to *k*-mer-based indexes built with PanKmer [4] and KMC3 [11]. PanKmer is a tool for reference-free pangenome analysis and stores presence/absence values of all 31-mers across the total genome collection for each genome [4]. KMC3 is a more generic *k*-mer counting tool and is very efficient in practice. We adapted KMC3 for the pangenome membership and conservation queries as described in Methods 2.6. We abbreviate the KMC3 membership and conservation queries as KMC3-M and KMC3-C, respectively. Likewise, we abbreviate the MEMO’s membership and conservation queries as MEMO-M and MEMO-C. We also indexed and evaluated MEMO-C for approximate conservation counts to the nearest decile threshold and refer to this as MEMO-DC (“DC” standing for “decile conservation”). We refer to PanKmer’s conservation query as PanKmer; PanKmer cannot perform the membership query.

We compared how these methods scale to two pangenomes: a human pangenome and a vertebrate pangenome. The human pangenome is composed of the autosomal chromosomes from 88 haplotypes from the Human Pangenome Reference Consortium (HPRC) and T2T-CHM13 [27, 19]. We refer to this collection as the HPRC pangenome, even though T2T-CHM13 is not part of the HPRC Year 1 data freeze release. We refer to the vertebrate pangenome, composed of 16 high-quality vertebrate genomes from the Vertebrate Genomes Project’s initial release, as the VGP pangenome [20]. Finally, we visualized sequence conservation from MEMO-C output across the human leukocyte antigen (HLA) locus of the HPRC pangenome as anchored to T2T-CHM13.

### 3.1 Indexing

The MEMO indexes were substantially smaller than equivalent *k*-mer-based indexes. The MEMO index for the HPRC pangenome, using T2T-CHM13 as the pivot genome, was roughly 2 GB. The MEMO-M index was by far the smallest: 539.2x smaller than the equivalent KMC3-M index. The MEMO-C index was 11.4x and 8.8x smaller than the equivalent PanKmer and KMC3-C indexes respectively (Table 1).

**Table 1.** Index and query statistics of pangenome query tools. The pangenome includes 88 human autosomal haplotypes from HPRC and T2T-CHM13. Index query types include: 1. Global presence/absence; 2. Member presence/absence; 3. Conservation; 4. Decile conservation. Query type 4* indicates no relative size reduction in a KMC3 decile index. The decile conservation index yields counts to the nearest lowest decile. Elapsed conservation query runtime and peak memory usage on the HLA locus (chr6:29476949-33231258) anchored to T2T-CHM13. Time is expressed in hours:minutes:seconds.

| Method | Index - HPRC |  |  |  | Query - HLA Locus |  |
| --- | --- | --- | --- | --- | --- | --- |
|  | Size (GB) | Pivot | Query Length | Query Type | Time | Memory (GB) |
| PanKmer | 23.29 | any | 31-mer only | 1, -, 3, - | 1:24:33.87 | 6.27 |
| KMC3-M | 1,267.20 | any | re-index | 1, 2, 3, - | 1:31:23.07 | 14.32 |
| KMC3-C | 18.05 | any | re-index | 1, -, 3, 4* | 0:00:35.71 | 18.10 |
| MEMO-M | 2.35 | re-index | any | 1, 2, 3, - | 0:00:51.15 | 2.69 |
| MEMO-C | 2.04 | re-index | any | 1, -, 3, - | 0:00:13.89 | 2.79 |
| MEMO-DC | 0.87 | re-index | any | -, -, -, 4 | 0:00:08.12 | 2.46 |

We separately measured the size of the compressed files produced using the MEMO Parquet strategy versus the strategy of using block-based bgzip compression and tabix indexing [14]. Parquet compression using the ZSTD codec yielded index sizes roughly 4x smaller than those produced by bgzip and tabix (Table 2). Notably, indexing the overlapping intervals between consecutive MEMs and order-MEMs yielded a better compression ratio compared to indexing the MEM intervals themselves.

**Table 2.**
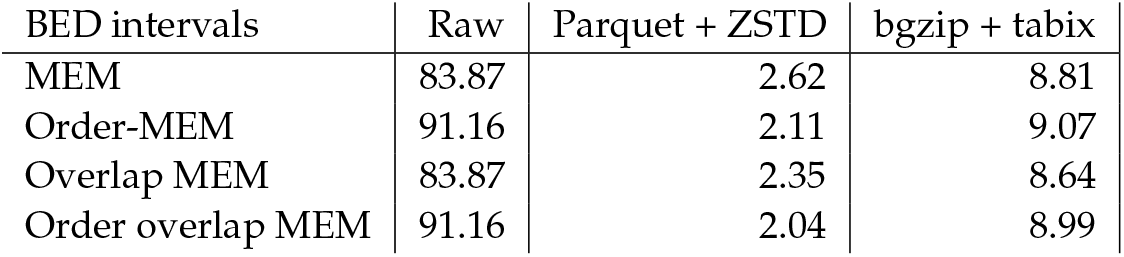
Comparison of approaches for compressing BED files (GB). MEMO-M is the compressed overlap MEM file. MEMO-C is the compressed order overlap MEM file. Parquet is used for columnar compression and file access, as compared to bgzip and tabix.

**Table 3.** HPRC index creation resources. Time (hours:minutes:seconds), CPU %, and peak memory usage as output from `/usr/bin/time`. PanKmer was run in 20 rounds with 8 threads and gzip-level 6. KMC3 was run with 8 threads and 20 GB max RAM. MEMO index creation relies on MONI to find MSs. MONI was run single-threaded during the build step and with 8 threads during MS finding.

| Method | Time | CPU % | Memory (GB) |
| --- | --- | --- | --- |
| PanKmer | 30:18:55 | 684% | 33.07 |
| KMC3-M | 03:40:20 | 514% | 20.03 |
| KMC3-C | 05:38:59 | 380% | 20.03 |
| MEMO-M | 23:13:49 | 4053% | 151.58 |
| MEMO-C | 24:38:07 | 3827% | 151.58 |

**Table 4.** VGP index resources. Time (hours:minutes:seconds), CPU %, and peak memory usage as output from `/usr/bin/time`. PanKmer was run in 20 rounds with 1 thread and gzip-level 6. KMC3 was run with 48 threads [default, max no. of CPU cores] and 12 GB max RAM. MEMO index creation relies on MONI to find MSs. MONI was run with 8 threads for the build and MS finding steps.

| Method | Time | CPU % | Memory (GB) |
| --- | --- | --- | --- |
| PanKmer | 15:48:00 | 99% | 273.52 |
| KMC3-M | 00:07:50 | 740% | 12.02 |
| KMC3-C | 01:27:15 | 173% | 12.03 |
| MEMO-M | 13:45:24 | 839% | 278.85 |
| MEMO-C | 14:37:56 | 799% | 278.85 |

**Table 5.** Index scalability to the HPRC pangenome. Index sizes in GB of an increasing number of HPRC autosomal haplotypes. MEMO indexes are anchored to T2T-CHM13.

| # Genomes | MEMO-M | MEMO-C | KMC3-M | KMC3-C | PanKmer |
| --- | --- | --- | --- | --- | --- |
| 9 | 0.33 | 0.34 | 128.19 | 15.41 | 14.57 |
| 18 | 0.59 | 0.59 | 256.31 | 15.85 | 15.58 |
| 27 | 0.81 | 0.80 | 384.42 | 16.21 | 16.54 |
| 36 | 0.97 | 0.95 | 512.58 | 16.48 | 17.39 |
| 45 | 1.16 | 1.13 | 654.98 | 16.79 | 18.36 |
| 54 | 1.36 | 1.31 | 768.88 | 17.11 | 19.36 |
| 63 | 1.57 | 1.51 | 897.07 | 17.43 | 20.40 |
| 72 | 1.94 | 1.74 | 1025.16 | 17.68 | 21.22 |
| 81 | 2.14 | 1.90 | 1153.31 | 17.87 | 22.44 |
| 89 | 2.35 | 2.04 | 1267.16 | 18.05 | 23.29 |

**Table 6.** Index scalability to the VGP pangenome. Index sizes in GB of an increasing number of VGP genomes. MEMO indexes are anchored to the blenny genome.

| # Genomes | MEMO-M | MEMO-C | KMC3-M | KMC3-C | PanKmer |
| --- | --- | --- | --- | --- | --- |
| 4 | 1.79 | 1.93 | 37.95 | 34.63 | 31.68 |
| 8 | 3.76 | 4.18 | 65.86 | 59.31 | 56.41 |
| 12 | 5.64 | 5.41 | 100.51 | 96.23 | 94.09 |
| 16 | 7.41 | 6.61 | 131.23 | 129.81 | 126.57 |

### 3.2 Pangenome scaling

MEMO enables approaches to reduce index sizes for large pangenomes. Although MEMO has a larger scaling factor than KMC3 and PanKmer for the HPRC pangenome, MEMO has comparable scaling to the VGP pangenome and can incorporate additional subsetting to reduce index size.

Across 9 to 89 HPRC haplotypes, MEMO index sizes roughly increase 6.5x, but are likely to remain under 4 GB for a large number of haplotypes. For the HPRC pangenome, KMC3 indexes scale 1.2x for KMC3-C and 9.9x for KMC3-M. A new KMC3 database must be made for each genome for the membership query; these together compose the KMC3-M index. PanKmer scales roughly 1.6x (Figure 2**A**). Across 4 to 16 vertebrate genomes from the VGP pangenome, MEMO indexes scale 3.8x; KMC3 indexes scale 3.6x; and PanKmer index scale 4.0x (Figure 2**B**). *K*-mer-based indexes scale poorly to diverse sets of genomes as a *k*-mer table must store each *k*-mer in the union of sequences. On the other hand, MEMO stores genome coordinates that are efficiently compressed.

**Figure 2.**
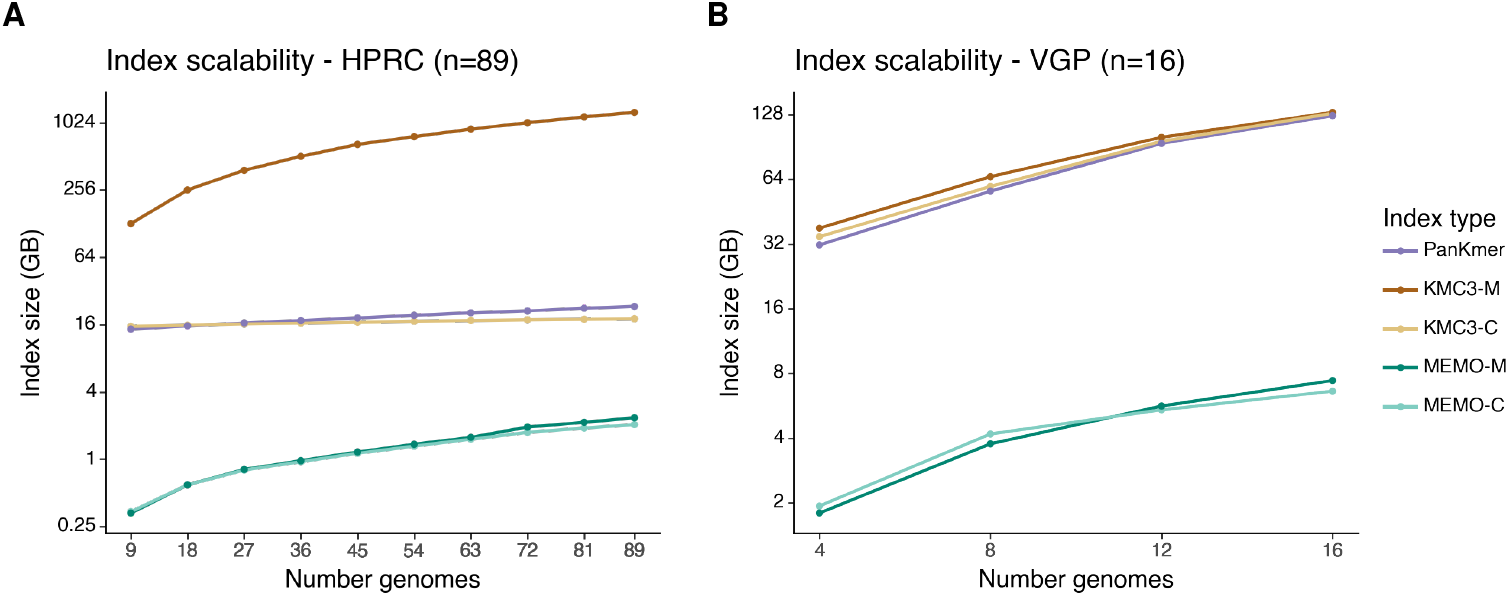
Index scalability of PanKmer, KMC3, and MEMO indexes. The X-axis is the number of indexed genomes. The Y-axis is the *log*2 index size (GB). **(A)** Index scalability across 89 autosomal HPRC haplotypes, anchored to T2T-CHM13. **(B)** Index scalability across 16 VGP genomes, anchored to the blenny genome.

The MEMO-C index sizes can further be reduced by leveraging the rank-ordered design to yield approximate conservation counts. Subsetting indexed orders to the deciles of 89 haplotypes, reduces the MEMO-C HPRC index to 0.87 GB (Table 1). Order subsetting allows the potential for small index sizes in larger pangenomes while still capturing pangenome sequence divergence. Subsetting the KMC3-C database to yield counts to the nearest decile yields no reduction in the index size since all *k*-mers must still be stored. PanKmer API does not have any functionality to reduce index size.

Limiting the permitted *k*-mer lengths for the queries allows further reduction in MEMO index size. Subsetting to overlap MEMs ≤ *t* restricts *k*-mer queries to *k < t*. In practice for the HPRC pangenome, restricting queries to *k* ≤ 31 (and so discarding MEMs length ≥ 32) reduces the number of MEM overlaps indexed by MEMO-M by 12.38% and the number of order-MEM overlaps indexed by MEMO-C by 18.55%. Removing these larger intervals allows for better compression and results in index sizes of 1.79 GB and 1.21 GB, respectively. Subsetting order and intervals allow MEMO two opportunities to reduce index size for larger pangenomes—approaches that are incompatible with *k*-mer-based indexes. The MEMO-DC HPRC index for conservation decile *k*-mer queries with *k* ≤ 31 is 0.67 GB.

### 3.3 Querying pangenome membership & conservation

MEMO queries are faster and more memory efficient than equivalent queries on *k*-mer-based indexes. MEMO queries 31-mer conservation across the human leukocyte antigen (HLA) locus on T2T-CHM13, a highly variable 3.75 Mbp region on Chromosome 6, in 13.89 seconds—2.6x and 365.3x faster than KMC3-C and PanKmer. KMC3-C and PanKmer peak memory usage is 5.1x and 2.2x more than MEMO-C (Table 1). The HRPC decile conservation MEMO-DC index exhibits further query speed and memory savings. Compared to KMC3-M, MEMO-M is 107.2x faster and uses 5.3x less peak memory. As the MEMO query runtime is proportional to the number of overlap MEM intervals, the runtime is roughly constant across varying-*k* for the same query region. On the other hand, to vary the length-*k k*-mer query, KMC3 indexes require re-indexing. PanKmer can only index 31-mers.

MEMO allows exploring visualizations of sequence conservation from varying-*k*. From MEMO-C, we visualized 31-mer conservation of the HLA locus of T2T-CHM13 across the HPRC pangenome (Figure 3), as inspired by Panagram [9]. The HLA locus sequence conservation plot captures known regions of high single nucleotide polymorphism density [25, 12]. Zooming onto the HLA *delta* block, we found that the conservation decile count approximation of MEMO-DC yields a similar sequence conservation plot as the full MEMO-C resolution, yet with a 2.3x smaller index. While conservation plots can be generated from KMC3-C and PanKmer, visualizations made using these tools will generally be limited to a fixed value of *k*. Their slower query speeds restrict practical interactive exploration, while MEMO-C’s faster query times allow interactive visualization and exploration.

**Figure 3.**
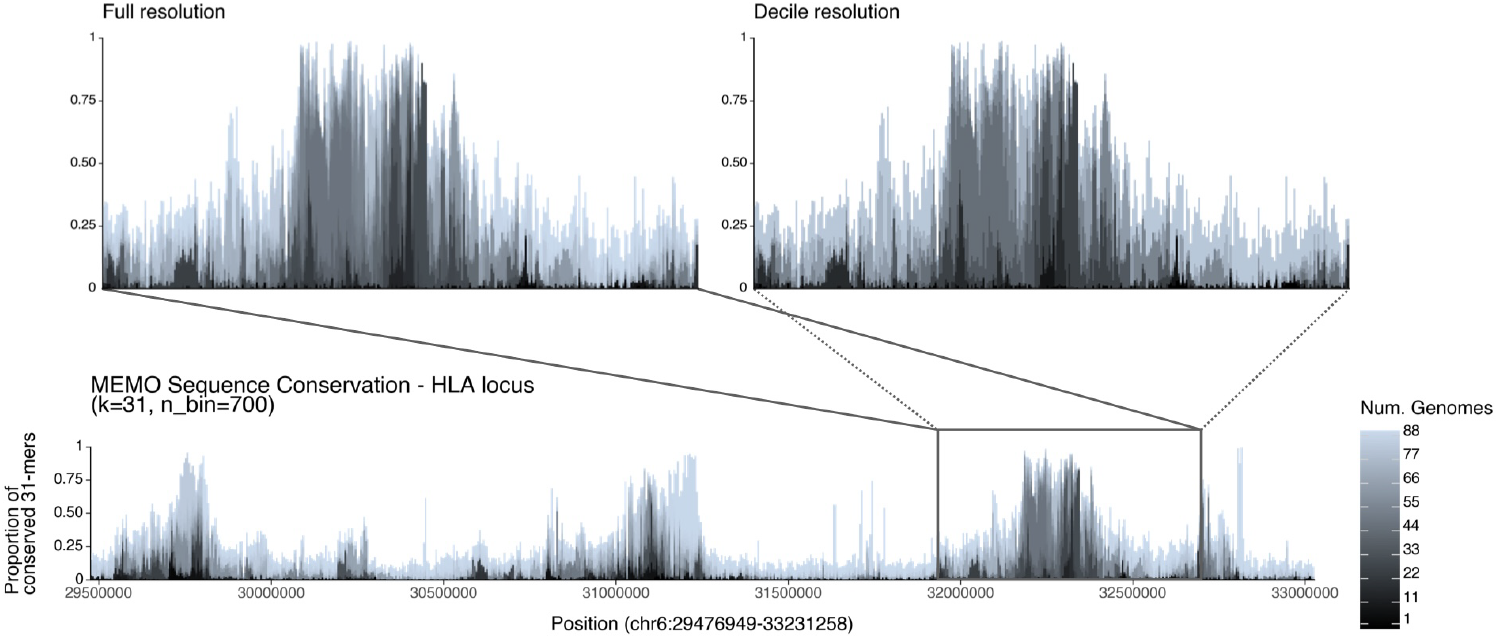
Sequence conservation plot from 31-mers anchored to the T2T-CHM13 HLA locus across the HPRC haplotypes (n=88). The user specifies a target region on the pivot, a length-*k*, and a histogram bin count to visualize the proportion of genomes containing the *k*-mer at each position of the query. The white area above the stacked bars represents the proportion of *k*-mers found across all 89 genomes. (Top left) Zoomed-in on the HLA *delta* block highlights a region of low sequence diversity. (Top right) Decile resolution of the *Delta* block demonstrates that MEMO-DC yields a plot that’s largely indistinguishable from the full-fidelity plot made by MEMO-C.

## 4 Discussion

We developed MEMO, a small MEM-based pangenome index that efficiently answers arbitrary-length *k*-mer membership and conservation queries. By using matching statistics as the basis for finding MEMs, we derived the related notion of order-MEMs, which are derived from matching statistics that have first been sorted across genomes. These ideas effectively generalize MEMs and matching statistics to the pangenomic context while enabling extremely small indexes.

MEMO’s fast query speed enables visual exploration of sequence conservation, especially in complex regions where the freedom to vary the *k*-mer length used can help to better understand distinct patterns of sequence conservation.

Indexing the overlapping intervals between consecutive MEMs and order-MEMs yielded a better compression ratio compared to indexing the MEMs themselves. Columnar compression using Parquet and ZSTD yielded roughly 4x better compression than commonly used bgzip and tabix. These observation could have wider significance in bioinformatics; switching to columnar compression may yield improved compression in other contexts.

MEMO’s chief limitation is the fact that a single pivot genome must be selected at index construction time. Although pangenomes typically do have a natural pivot—i.e. a genome that has a higher quality assembly or annotation compared to the others—there could also be situations where no natural pivot exists. The VGP project is an example of this. In the future, it will be important to consider designing multi-pivot generalizations of MEMO, which could possibly benefit even more from the inherent redundancy of the pangenome.

While MEMO uses MONI to find matching statistics, MONI is not tailored to our problem. Instead, the profile document array of Ahmed et al. could be used in the future [3]. MEMO demonstrates the potential of MEM-based indexes over *k*-mer-based indexes for compressed indexes and fast flexible queries on large pangenomes.

## 2012 ACM Subject Classification

Applied computing → Computational genomics

## Supplementary Material

The MEMO open-source software is available at https://github.com/StephenHwang/MEMO. The code used to run the experiments is available at https://github.com/StephenHwang/MEMO_experiments

## Funding

This work was carried out at the Advanced Research Computing at Hopkins (ARCH) core facility (rockfish.jhu.edu), supported by the National Science Foundation (NSF) grant OAC 1920103.

*Stephen Hwang*: Johns Hopkins University, XDBio Program

*Nathaniel K. Brown*: Johns Hopkins University Computer Science PhD Fellowship

*Omar Y. Ahmed*: R01HG011392 and R35GM139602 to BL

*Katharine M. Jenike*: NSF grant 2216612, NIH grant U01CA253481, and HFSP award RGP0025/2021 to MCS

*Sam Kovaka*: NSF grant 2216612, NIH grant U01CA253481, and HFSP award RGP0025/2021 to MCS *Michael C. Schatz*: NSF grant 2216612, NIH grant U01CA253481, and HFSP award RGP0025/2021 to MCS

*Ben Langmead*: R01HG011392 and R35GM139602 to BL

## Acknowledgements

We thank Christina Boucher for helpful conversations.

## A Appendix

